# Mitochondrial genomes of three *Melampus* species (Ellobiidae; Gastropoda) and a proposed mitochondrial control region for Euthyneura

**DOI:** 10.1101/2024.07.03.601860

**Authors:** Thomas Inäbnit, Ralph Tiedemann, Alice B. Dennis

## Abstract

*Melampus bidentatus* was recently divided into three genetically divergent cryptic species: *M. bidentatus, M. jaumei* and *M*. gundlachi. We have assembled and annotated a mitochondrial genome for each of these species. Comparisons with other taxa showed that these three species possess a gene order that is distinct from the conserved gene order found in the six (out of nine available) Ellobiid genera. Among the species with a derived gene order, mean nucleotide divergence over all genes was on average 1.57× higher than among mitogenomes with the conserved gene order. This suggests varying evolutionary rates of mitochondrial genes in this family. We also use comparisons among 130 mitochondrial genomes within Euthyneura to support the presence of a control region preceding the Isoleucin-tRNA, including in the novel genomes presented here.

## Introduction

The coffee-bean snails *Melampus bidentatus* Say 1822, *Melampus jaumei* Mittre 1841, and *Melampus gundlachi* Pfeiffer 1853 are cryptic air-breathing Ellobiids that were until recently considered conspecific (Dennis & Hellberg 2010, Inäbnit et al. 2023; Figure 1). They are distributed in distinct ranges: *M. bidentatus* occurs from New Brunswick to Florida, *M. jaumei* in the northern Gulf of Mexico, Atlantic Florida, and from North Carolina to Delaware, and *M. gundlachi* in southern Florida, southern Texas, and the Greater Antilles (Dennis & Hellberg 2010). The ranges are parapatric, with *M. bidentatus* and *M. jaumei* co-occurring between North Carolina and Delaware.

**Figure 1:**
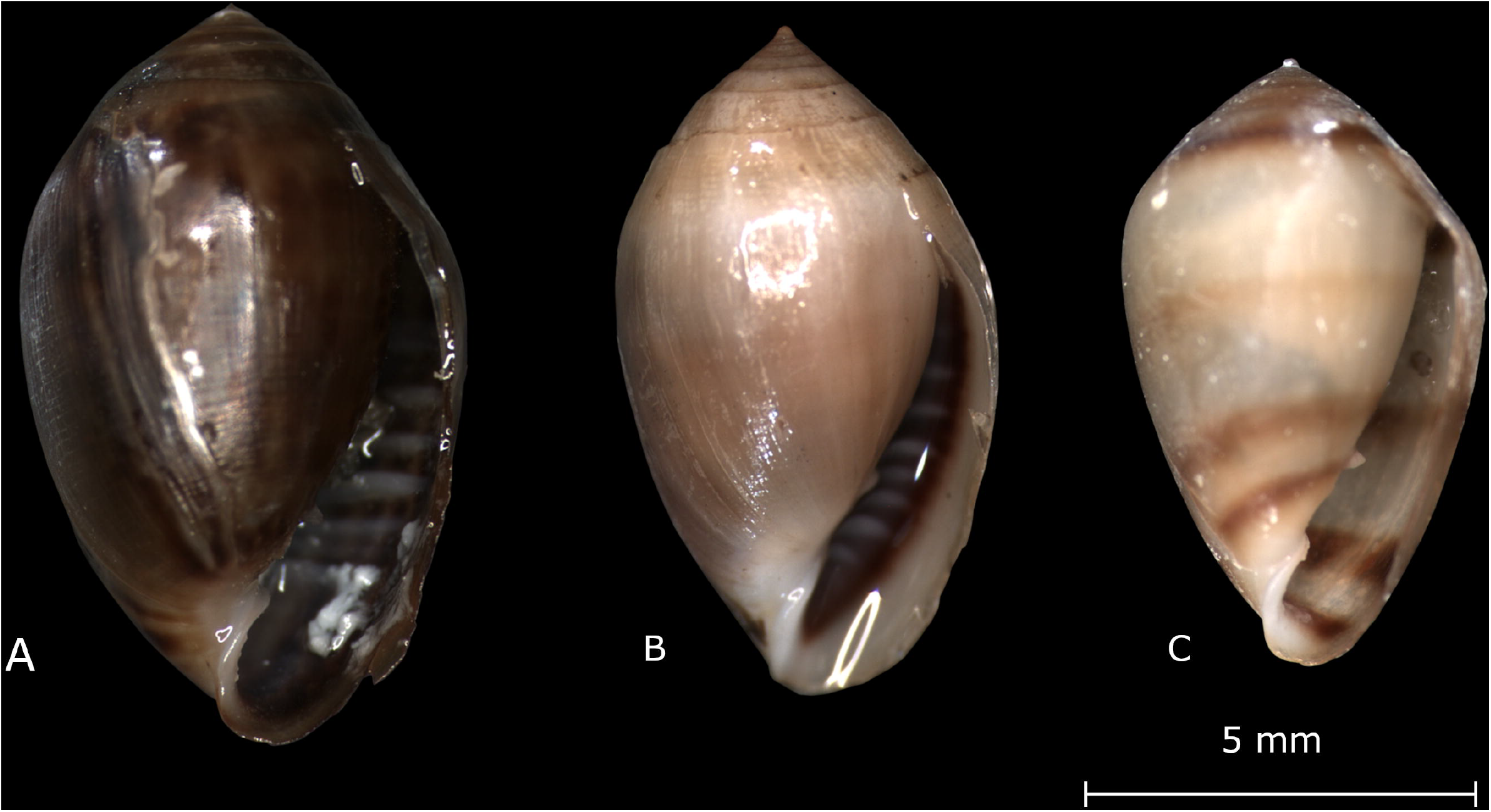
Images of the three *Melampus* species sequenced in this paper. A: *Melampus bidentatus* from Beebe Cove, Connecticut; B: *Melampus jaumei*, a specimen from the same collection site (Dauphin Island, Mobile Bay, Alabama) as the sequenced specimen; C: *Melampus gundlachi* from the same collection site (South Padre Island, Texas) as the sequenced specimen. Photos taken by Alice Dennis.

Analysis of COI sequences has shown high divergence between the species (22–24%; Dennis & Hellberg 2010), but other mitochondrial genes have not been analyzed. In other Ellobiid species, phylogenies constructed from whole mitogenomes (e.g. Varney et al. 2021) have recovered species which were assigned to the family based on other genetic markers (Romero et al. 2016a) and morphology (Martins 1995, 1996, 2007) outside the family (e.g. *Myosotella myosotis, Pedipes pedipes*). These ellobiids possess distinct mitochondrial gene orders relative to the conserved gene order within Euthyneura (i.e. Heterobranchia without “lower” taxa: White et al. 2011, Varney et al. 2021) and most ellobiids. This study aims to unravel the divergence across entire mitgenomes in the formerly cryptic *Melampus* species, and to place it in context with the mitogenome architecture across Euthyneura.

## Material & Methods

### Collections and sequencing

Three species were sequenced in this study, all collected in sites of allopatric occurrence (Table 1). For *M. bidentatus* DNA extraction and library preparation was performed by Dovetail Genomics (Scotts Valley, CA, USA) with the TruSeq DNA PCR-Free kit (Trueq Index 5, Illumina, San Diego, CA, USA, mean size 475bp). 150 bp paired-end sequencing was performed on one lane of HiSeqX by Novogene (Cambridge, UK).

**Table 1:** *Collection sites for the three species for which mitochondrial genomes are presented here. Coverage in* M. bidentatus *was calculated by NOVOPlasty and refers to mitochondrial sequences only. For the other two species, the coverage was calculated by comparing the number of unassembled PacBio HiFi-reads with the number of contigs in the MitoFinder assembly*.

| **Species** | **Collection site** | **Collection date** | **Coverage** |
| --- | --- | --- | --- |
| *Melampus bidentatus* | Barn Island,<br>Connecticut<br>(Latitude: 41.340,<br>Longitude: -71.875) | 11. October 2017 | 125× |
| *Melampus jaumei* | Mobile Bay, Alabama<br>(Lat.: 30.251,<br>Long.: -88.201) | 8. December 2020 | 24.72× |
| *Melampus gundlachi* | South Padre Island,<br>Texas, USA<br>(Lat.: 26.139,<br>Long.: -97.176) | 16. May 2006 | 63.23× |

For *Melampus jaumei*, DNA was extracted using a modified CTAB protocol (Toonen 2001). Extracted DNA was sent to Novogene (Cambridge, UK) for library preparation and PacBio HiFi sequencing.

For *Melampus gundlachi* DNA was extracted using the MagAttract HMW kit (Qiagen 67536) tissue protocol, and included an overnight tissue digestion. This DNA was sent to the University of Leiden for library preparation (low-input sequencing library) and PacBio HiFi sequencing.

Raw sequencing data has been deposited in NCBI Sequence Read Archives (*M. bidentatus*: SRR8415687; *M. jaumei*: SRR24965506; *M. gundlachi*: SRR24958157). The individuals sequenced in this study were destructively sampled during DNA extraction, but additional individuals from the same collection or locale have been deposited at the Florida Museum of Natural History (*M. jaumei*: 3 specimens, UFID 580985-7; *M. gundlachi*: 1 specimen, UFID 580978) and at the Royal Belgian Institute of Natural Sciences (*M. bidentatus*: 1 specimen, INV.302.000).

### Assembly, and annotation

From Illumina sequences (*M. bidentatus*), sequencing primers and low quality sequences were removed with Trimmomatic (Bolger et al. 2014) (minimum phred score of 15 over a 4bp sliding window). Bacterial, archeal, and viral contaminants were removed using KRAKEN (v minikraken_20171019_8GB; Wood & Salzberg 2014).

Illumina short reads from *M. bidentatus* were assembled with NOVOPlasty v3.8.3 (Dierckxsens et al. 2016) using default settings and seeded with a CO1 sequence from the same site and species (GenBank accession: HM154306). Mitogenomes of *M. jaumei* and *M. gundlachi* were assembled from unassembled PacBio Hifi data using MitoFinder v1.4 (Allio et al. 2020) with standard settings. MitoHiFi v2.0 (Uliano-Silva et al. 2022) was then used to extract and circularize the resulting mitochondrial genome sequence. Annotation was performed using the online MITOS webserver (Bernt et al. 2013) with the “Invertebrate Mitochondrial” genetic code. Annotations were manually checked for the presence of start and stop codons and adjusted if they had been incorrectly annotated.

To locate a putative control region, all non-coding regions from complete mitochondrial genomes, both produced here and from further Ellobiidae available on GenBank (Table 2, as of 07/2021) were aligned and compared. Specifically, we searched for non-coding regions in the same positions in all of these mitochondrial genomes. The presence of those non-coding regions was further evaluated in 151 Heterobranch (mostly Euthyneuran) mitochondrial genomes downloaded from GenBank (Supplementary table 2).

**Table 2:**
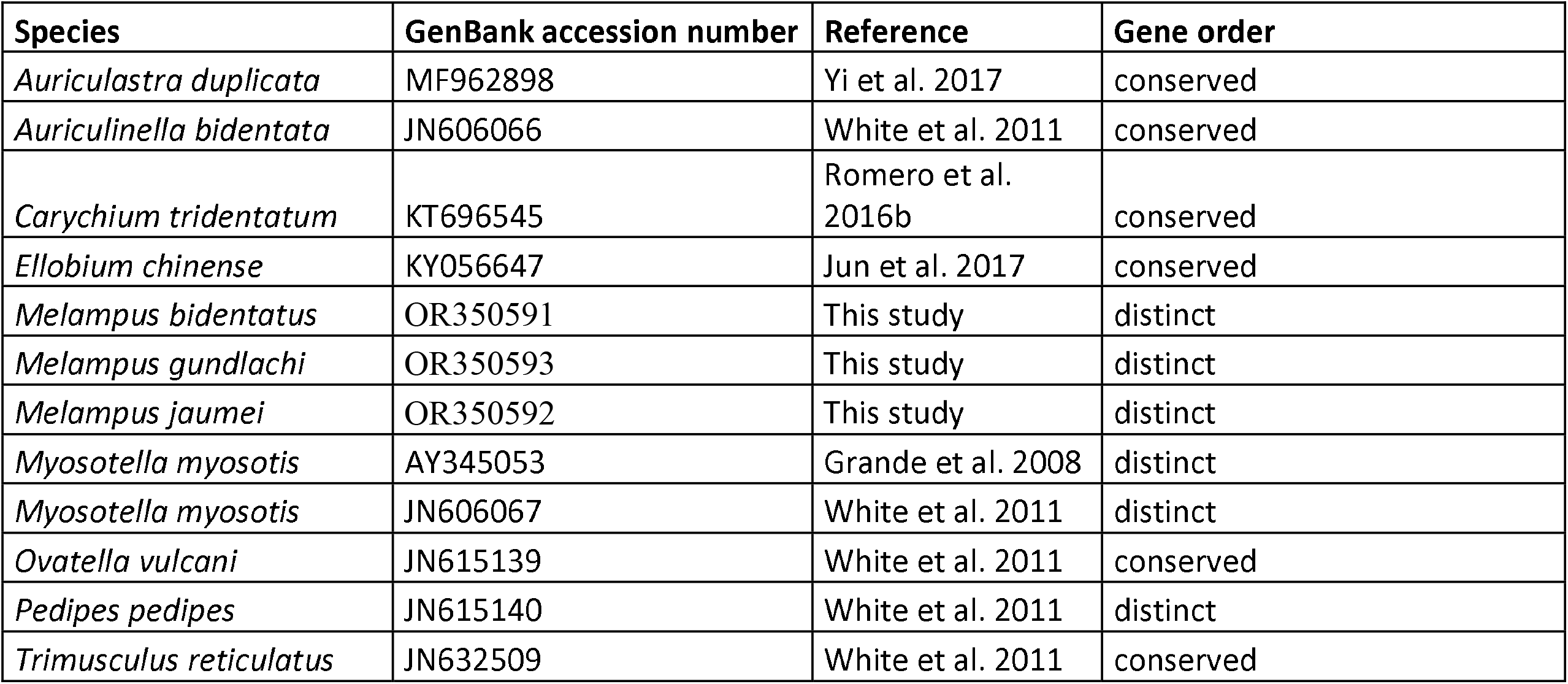
Mitochondrial genomes from Ellobiidae used in our analyses

All coding regions of all mitochondrial genomes within Ellobiidae (Table 2) were aligned using ClustalW as implemented in Mega X (Tamura et al. 2004, Kumar et al. 2018). Two separate subsets were created, containing (1) species with the conserved gene order of Euthyneura (White et al. 2011, Varney et al. 2021) and (2) species with derived gene orders. The overall mean distances were calculated for both subsets for each coding region and the amino acid sequence of all protein-coding genes, as well as for the full alignments of the three *Melampus* species and the ellobiids with conserved gene order in Mega X. For the three *Melampus* species, pairwise distances were calculated in Mega X for all genes separately and the mitochondrial genome as a whole.

Phylogenetic trees were calculated using the protein coding genes of all published ellobiid mitochondrial genomes (table 2; as of the end of 2023), as well as *Onchidella celtica* (Cuvier 1816; GenBank accession number AY345048) as the outgroup. Sequences were aligned using MAFFT v7.450 (Katoh et al. 2002; Katoh & Standley 2013) for Windows with the FFT-NS-2 algorithm. A maximum likelihood tree was calculated in IQTree 2.1.2 (Minh et al. 2020, Chernomor et al. 2016) with the ModelFinder function (Kalyaanamoorthy et al. 2017) and 1000 bootstrap replicates. A Bayesian tree was calculated in MrBayes 3.2.7 (Ronquist et al. 2012) using the mixed substitution model, invgamma rate variation and a Markov Chain Monte Carlo (MCMC) chain length of 10 000 000 generations and a subsampling frequency of every 4000 generations with the first 100 000 generations being discarded as burn-in, four heated chains and a chain temperature parameter of 0.2. The species tree of Romero et al (2016a) was redrawn in simplified form to facilitate comparisons with the mitochondrial genome trees presented here.

## Results & Discussion

The mitochondrial genomes of *M. bidentatus* (14668 bp; coverage: 125×), *M. jaumei* (14811 bp; coverage: 24.72×) and *M. gundlachi* (14823 bp; coverage: 63.23×) are identical in gene order (Fig. 2), similar in length, and contain 37 genes: 13 protein coding genes, 2 rRNAs and 22 tRNAs. Twenty four genes are encoded on the plus strand and 13 on the minus strand. Their gene order is distinct from the conserved gene order of Euthyneura and all genes encoded on the plus strand are grouped together, as are the genes encoded on the minus strand (Supplementary Table 1). The large divergence in COI sequences between the three species (Dennis & Hellberg 2010) was also present in all other mitochondrial genes in the mitochondrial genome (entire genome: 0.307-0.335, Supplementary table 3). The overall mean distance between our three cryptic *Melampus* species (0.32) is similar to distances among all ellobiid mitochondrial genomes with the conserved gene order (0.38, Supplementary table 4).

**Figure 2:**
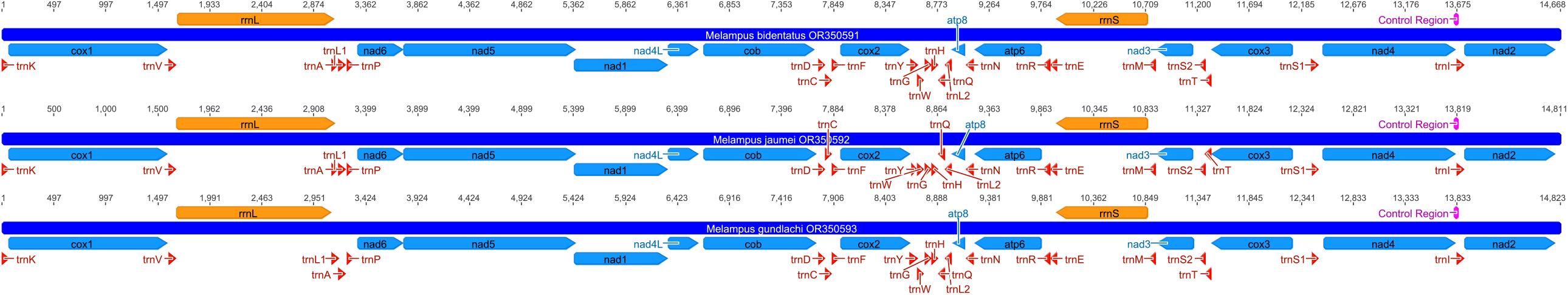
Mitochondrial genome map of *M. bidentatus, M. jaumei* and *M. gundlachi*. The map was drawn in Geneious v.8.0.5 (www.geneious.com).

The only non-coding mitochondrial region shared between all Ellobiidae is a short region preceding the Isoleucin tRNA (trnI). Its length is similar in the three *Melampus* species (*M. bidentatus*: 12 bp; *M. jaumei*: 11 bp; *M. gundlachi*: 13 bp) and more variable across Ellobiidae (11-317 bp, Supplementary table 2). This non-coding region is present in at least 130 of the 151 analyzed mitochondrial genomes within Euthyneura (Supplementary table 2; many of the remaining 21 mitochondrial genomes had gene order rearrangements in that region). This stretch has been previously suggested to comprise the control region by other studies using much smaller datasets (Grande et al. 2008, White et al. 2011, Kurabayashi & Ueshima 2000) and our study shows it to be widespread across mitochondrial genomes of Euthyneura.

In the phylogenetic trees based on the mitochondrial protein-coding genes (Fig. 3A) the three *Melampus* species cluster with the other ellobiid species (*Pedipes pedipes* and *Myosotella myosotis*) with a derived mitochondrial gene order (Bayesian Posterior Probabilities (BPP): 0.985; Bootstrap Support (BS): 46%). This group is separated from the group containing the species with a conserved gene order. Among the species with derived gene order, *P. pedipes* and *M. myosotis* form a decently supported (BS: 71%; BPP: 0.921) clade. The position of *Ovatella vulcani* differs between the Maximum Likelihood (grouping outside a group consisting of *Ellobium chinense, Auriculastra duplicata & Auriculinella bidentate*; BS: 33%) and Bayesian trees (sister group of *Carychium tridentatus*; BPP: 0.797), but neither pairing has high node support values. In the most current published phylogeny (Romero et al. 2016a, fig. 1; redrawn here in a simplified form as Fig. 3B), *M. myosotis* is instead recovered as the sister species of *O. vulcani*, a species with conserved gene order, which is consistent with the classification of both species within the subfamily Pythiinae in morphological studies (e.g. Martins 2007). Romero et al. (2016a) also recovered *Trimusculus reticulatus* as the sister group of all other ellobiids, including those with derived mitochondrial gene orders which were recovered separately in the mitochondrial phylogeny (Fig. 3A).

**Figure 3:**
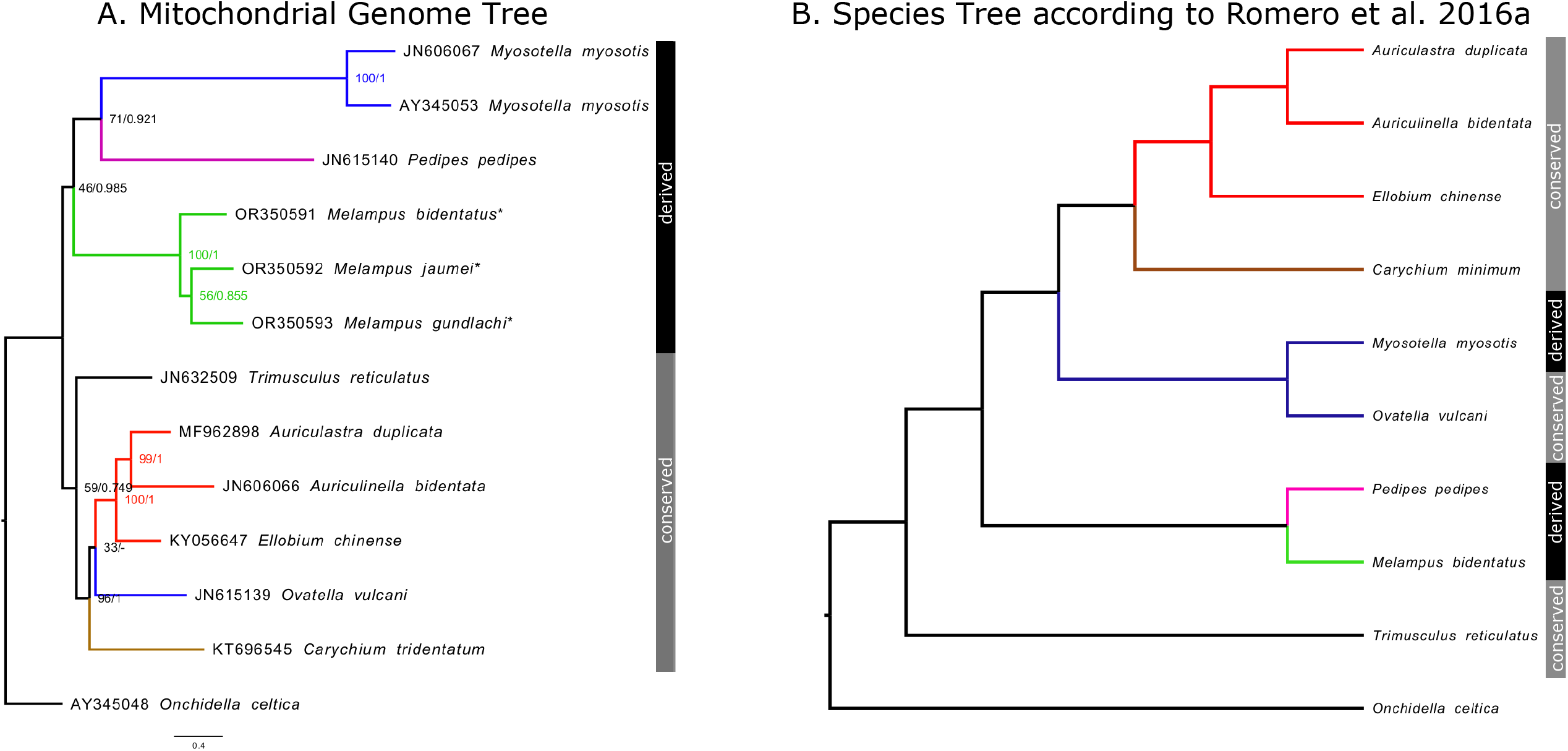
A) Phylogenetic Tree of all published ellobiid mitochondrial genomes. Mitochondrial genomes sequenced in this study are marked with a star (*). Node support values of both the Maximum Likelihood analysis (left) and the Bayesian Inference (right) are given. B) Simplified redrawing of Romero et al. (2016a), fig.1 using the same or related species as in A). Branch color represents subfamilies: green: Melampodinae, purple: Pedipediinae, brown: Carychiinae, red: Ellobiinae, blue: Pythiinae, black: Trimusculiinae and the outgroup (*Onchidella celtica*). The black (conserved gene order) and grey (derived gene orders) bars to the right of the tree indicate whether a species has the conserved euthyneuran mitochondrial gene order or a derived mitochondrial gene order.

Among Ellobiidae, the overall mean distance among mitochondrial genes is (except for three tRNAs) consistently higher (on average 1.57×) in taxa with a distinct gene order than among those with the conserved gene order (Supplementary table 4). This difference in divergence between conserved and distinct gene orders might indicate that the evolutionary rates of mitochondrial genomes vary within Ellobiidae (varying evolutionary rates in ellobiid mitochondrial genes have also been observed by Inoue et al. 2022). This can lead to false groupings of taxa with accelerated evolutionary rates in phylogenetic analyses (Liu et al. 2018), which might explain why ellobiid mitochondrial genomes with distinct gene orders are generally recovered outside Ellobiidae in Panpulmonate or Heterobranch phylogenies (such as Varney et al. 2021).

In conclusion, our new mitogenomes have shown that the deep divergence previously documented in a single mitochondrial gene for *Melampus* (CO1, Dennis & Hellberg) extend to all coding genes in the mitochondria. These new mitogenomes provide a resource for further studies within the genus *Melampus*, across ellobids, and add to our broad understanding of evolutionary rates in mitochondrial genes. Lastly, we provide further support to the suspected position of the control region within Euthyneura mitogenomes (Grande et al. 2008, White et al. 2011, Kurabayashi & Ueshima 2000).

## Acknowledgments

We thank Steven Loomis for collecting *M. bidentatus* and Morgan Kelley for collecting *M. jaumei*. Calculations were performed on the hpc-cluster of the ZIM at the University of Potsdam.

## Funding

This project was financed with a DFG grant awarded to A. Dennis (DE 3002/3-1). Sequencing of *M. bidentatus* was supported by an Academic Transition Grant awarded to A. Dennis by EAWAG (Swiss Federal Institute of Aquatic Science and Technology).

## Conflict of Interest

The authors do not have a conflicts of interest to declare.

## Authors contributions

T.I. and A.B.D. designed the research with input from R.T. A.B.D. processed the samples in the laboratory and pre-processed Illumina short-read data. T.I. assembled and analyzed the mitochondrial genomes with input from A.B.D. and R.T. T.I. wrote the manuscript with the help of A.B.D. and R.T..

## Notes

### Competing Interest Statement

The authors have declared no competing interest.

